# Gene co-expression reveals the modularity and integration of C_4_ and CAM in *Portulaca*

**DOI:** 10.1101/2021.07.07.451465

**Authors:** Ian S. Gilman, Jose J. Moreno-Villena, Zachary R. Lewis, Eric W. Goolsby, Erika J. Edwards

## Abstract

C_4_ photosynthesis and Crassulacean acid metabolism (CAM) have been considered as largely independent adaptations in spite of sharing key biochemical modules. *Portulaca* is a geographically widespread clade of over 100 annual and perennial angiosperm species that primarily use C_4_, but facultatively exhibit CAM when drought stressed, a photosynthetic system known as C_4_+CAM. It has been hypothesized that C_4_+CAM is rare because of pleiotropic constraints, but these have not been deeply explored. We generated a chromosome-level genome assembly of *P. amilis* and sampled mRNA from *P. amilis* and *P. oleracea* during CAM induction. Gene co-expression network analyses identified C_4_ and CAM gene modules shared and unique to both *Portulaca* species. A conserved CAM module linked phosphoenolpyruvate carboxylase (PEPC) to starch turnover during the day-night transition and was enriched in circadian clock regulatory motifs in the *P. amilis* genome. Preservation of this co-expression module regardless of water status suggests that *Portulaca* constitutively operate a weak CAM cycle that is transcriptionally and post-transcriptionally upregulated during drought. C_4_ and CAM mostly used mutually exclusive genes for primary carbon fixation and it is likely that nocturnal CAM malate stores are shuttled into diurnal C_4_ decarboxylation pathways, but we find evidence that metabolite cycling may occur at low levels. C_4_ likely evolved in *Portulaca* through co-option of redundant genes and integration of the diurnal portion of CAM. Thus, the ancestral CAM system did not strongly constrain C_4_ evolution because photosynthetic gene networks are not co-regulated for both daytime and nighttime functions.

## Main

Modularity and evolvability have become central research areas in biology over the past three decades^1, 2^. Modularization generates simple building blocks that facilitate rapid and complex adaptations through the combination of more discrete functions and the ability of simpler units to explore mutational space with minimal pleiotropic effects^1–5^. Modern “-omics” strategies have enabled detailed resolution of gene regulation in diverse study systems, and modeling results have shown that carbon metabolic networks have high potential for exaptation into evolutionary innovations in bacteria^6^. Empirical research has identified deeply conserved carbon metabolic modules from anaplerotic pathways that have been repeatedly recruited into plant photosynthetic adaptations known as carbon concentrating mechanisms (CCMs)^7^. CCMs provide classic examples of rapid, adaptive evolution through the exaptation of existing metabolic modules that are subsequently refined for new functions.

Since the early Oligocene, CCMs—typically taking the form of C_4_ photosynthesis or Crassulacean acid metabolism (CAM)—have evolved many dozens, if not hundreds, of times independently^8–12^. Certain facets of CCMs, such as the main biochemical pathways, are evolutionarily accessible to all green plant lineages because they belong to deeply conserved photosynthetic and respiratory gene networks^7, 13^. It is thought that by coupling these gene networks to light responses (C_4_)^14^ and circadian oscillators (CAM)^15^, CCMs create two-stage carbon fixation pathways that address the most fundamental ecophysiological tradeoff in land plants: balancing CO_2_ influx with water loss to transpiration.

Most plants assimilate carbon using C_3_ photosynthesis, in which CO_2_ is directly fixed by RuBisCO during the day in mesophyll cells. However, RuBisCO can also bind to O_2_, triggering a set of costly reactions called photorespiration, which is exacerbated by hot, dry conditions^16^. CCMs increase the efficiency of photosynthesis by increasing the concentration of CO_2_ around RuBisCO. This is achieved by first fixing carbon with phosphoenolpyruvate (PEP) carboxylase (PEPC), which is either spatially (C_4_) or temporally (CAM) decoupled from final fixation of CO_2_ by RuBisCO (Fig. 1a, b). In both CCMs, CO_2_ is first captured by beta carbonic anhydrase (BCA) and PEPC, forming oxaloacetate (OAA) (Fig. 1a, b) in the mesophyll, but exclusively during the night in CAM (Fig. 1b). OAA is then reduced to malate by malate dehydrogenase (MDH) and stored in the vacuole overnight as malic acid. During the day, CAM plants limit gas exchange with the external environment, and instead decarboxylate stored malate with malic enzymes (MEs) to release CO_2_. Most C_4_ species also convert OAA to malate (NADP-type biochemistry), but others first transform it to aspartate (NAD-type biochemistry) via aspartate aminotransferase (ASP). These 4-carbon acids are transported to the bundle sheath cells and decarboxylated. In NAD-type C_4_ species, ASP reversibly transforms aspartate back into OAA in bundle sheath mitochondria, where it is reduced to form malate by NAD-MDH and finally decarboxylated by NAD-ME. By separating initial CO_2_ capture in the mesophyll from assimilation in the bundle sheath, C_4_ increases photosynthetic rates and water use efficiency while minimizing energy and carbon lost to photorespiration^17–19^. CAM does not always reduce photorespiration^20, 21^, but greatly increases water use efficiency^22, 23^ and reduces the harmful effects of high heat and radiation caused by oxidative stress^20^.

**Figure 1.**
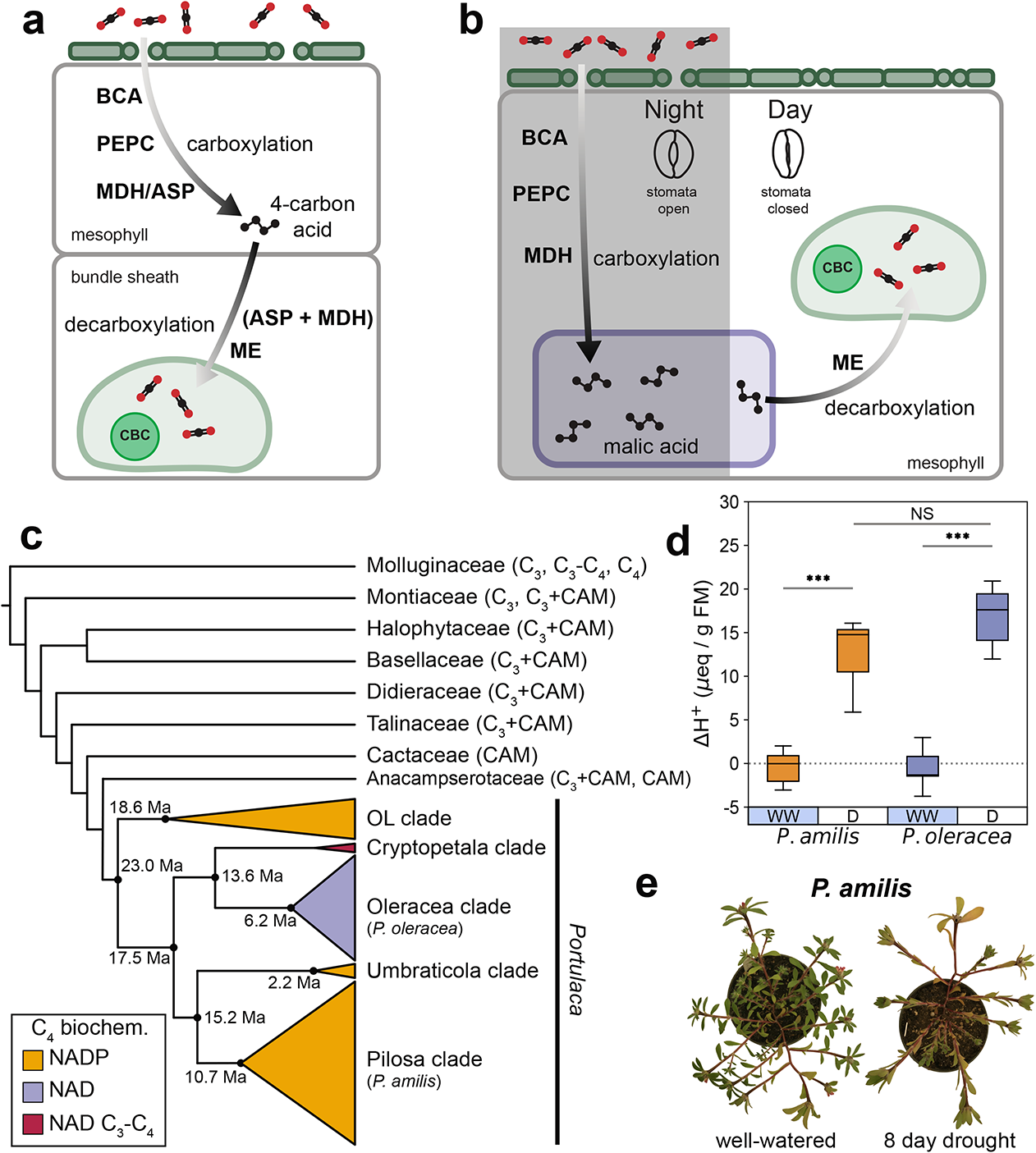
Simplified C_4_ (a) and CAM (b) pathways showing shared carboxylation and decarboxylation pathways. ASP, aspartate aminotransferase; BCA, beta carbonic anhydrase; CBC, Calvin-Benson cycle; MDH, malate dehydrogenase; ME, malic enzyme; PEPC, PEP carboxylase. Cartoon dendrogram of the Portullugo (Portulacineae + Molluginaceae), highlighting major *Portulaca* clades (c), which are colored by primary C_4_ biochemical pathway. Mean age estimates for focal *Portulaca* clades are from Ocampo and Columbus^56^. Night-day differences in titratable acidity of experimental *P. amilis* (orange) and *P. oleracea* plants (purple) (d). WW, well-watered; D, drought. Asterisks connecting lines indicate significant differences between well-watered and drought treatments (independent *t-*test; “***”, *p* < 0.001; “NS”, *p* = 0.121). Images of *P. amilis* when well-watered and after 8 days without water (e).

C_4_ and CAM boost the efficiency of photosynthesis in different ways and are therefore hypothesized to have evolved in response to different stressors. C_4_ has been primarily considered as an adaptation to high rates of photorespiration^24, 25^, and CAM to water limitation^8, 26^, though photorespiratory stress and drought stress are often experienced simultaneously^27^. Given the overlapping environmental contexts of CAM and C_4_ evolution, their repeated assembly of a similar set of biochemical modules^28^ (Fig. 1a, b), and their apparent ease of evolution, it is perhaps surprising that use of both C_4_ and CAM (hereafter C_4_+CAM) has only been reported in three lineages: *Portulaca* (Portulacaceae)^29^, *Spinifex* (Poaceae)^30^, and *Trianthema* (Aizoaceae)^31^. On the other hand, it may be precisely the large overlap in CCM gene networks and metabolites that may make C_4_ and CAM mutually exclusive in most lineages^32^. Plants that use both C_4_ and CAM seemingly must regulate the same gene networks in contrasting patterns that may result in futile metabolite cycling and inefficient transport. For example, daytime decarboxylation activity in the mesophyll during CAM would directly oppose C_4_carboxylation activity.

*Portulaca*, a geographically widespread clade of over 100 annual and perennial herbs and small shrubs, predominantly use C_4_, but facultatively exhibit CAM in response to drought^28, 29, 33–41^. It has been hypothesized that their CAM cycle must be either spatially separated from the C_4_ cycle, that is, relegated to a subpopulation of leaf cells such as water storage or bundle sheath^32, 35^; or alternatively, that *Portulaca* operate a novel two cell CAM system, whereby CAM malate is decarboxylated in the bundle sheath^36, 37^. Facultative CAM, the reversible induction of CAM in response to drought stress, is assumed to be ancestral in *Portulaca* because it has been observed in every major sub-clade assayed^38, 39^ as well as in *Portulaca*’s closest relatives^42–47^ (Fig. 1c). Transcriptomic studies of *Portulaca* and its close relatives suggest that the ancestral CAM pathway used both NAD- and NADP-enzymes for CAM^40, 41, 44^. The biochemical diversity of C_4_ pathways in major *Portulaca* clades and the observation of intermediate photosynthetic phenotypes (C_3_-C_4_) in the Cryptopetala clade^39, 48, 49^ imply multiple origins of C_4_ within a facultative CAM context (Fig. 1c). Multiple C_4_ origins are further evidenced by differences in leaf ultrastructure among lineages^48–50^, as Kranz anatomy is typically established early in C_4_ evolution^25, 51^.

Photosynthetic gene activity has only been studied in detail in *P. oleracea*^40, 41^, an NAD-type species. Similar experiments on other *Portulaca* are needed to answer fundamental questions about the function and evolution of C_4_+CAM, and the integration of new metabolic modules more generally. Facultative CAM is hypothesized to be ancestral to *Portulaca*, but expression of CAM-specific orthologs in multiple species has not yet been documented. And although multiple carboxylation and decarboxylation pathways can co-exist in some C_4_ plants^52^, it is not obvious why or how different C_4_ biochemistries would evolve in lineages that shared an ancestral, mixed CAM biochemistry type, nor do we know the extent to which these CCMs have been integrated in various lineages.

Here, we present a chromosome-level genome assembly of *Portulaca amilis* (NADP-type C_4_), the first of any C_4_+CAM plant. We analyzed transcriptomic data during a CAM-induction experiment to understand the synergistic evolution of multiple metabolic modules. Concurrent transcriptomic data for *P. oleracea* allowed us to compare C_4_+CAM systems across a deep split in the *Portulaca* phylogeny and discriminate between ancestral, inherited and lineage specific gene networks. We confirm previous hypotheses that *Portulaca* species share one CAM-specific PEPC ortholog *(PPC-1E1c*) and another C_4_-specific ortholog *(PPC-1E1a’*)^28^, and that CAM evolved by linking PEPC to starch catabolism via the circadian clock^13^. We found some evidence for CAM-specific decarboxylation pathways, but the diurnal portion of CAM appears to have been largely integrated into C_4_ metabolism, and we predict that nocturnal acids produced by CAM are likely shuttled into the diurnal pool of bundle sheath C_4_-acids. Finally, although *P. amilis* and *P. oleracea* share some central C_4_ orthologs, exclusive use of orthologs from most core gene families highlights the diversity of C_4_+CAM systems and provides further evidence that C_4_ evolved largely independently in multiple *Portulaca* lineages.

## Results

### CAM induction experiment

Significant diel fluctuations in titratable leaf acidity indicative of CAM activity were observed after seven and eight days without water in *P. amilis* and *P. oleracea*, respectively (Fig. 1d). Diel fluctuations in titratable acidity were not significantly different from zero when well-watered (one sample *t*-test; *t_amilis_*(6) = −0.522, *p_amilis_* = 0.624; *t_oleracea_*(6) = −0.573, *p_amilis_* = 0.591), but were significantly different under drought (one sample *t*-test; *t_amilis_*(6) = 7.437, *p_amilis_* = 0.00069; *t_oleracea_*(6) = 10.136, *p_amilis_* = 0.00053), and significantly greater than when well-watered (Fig. 1d). There was no significant difference between the magnitude of diel fluctuations between species (independent *t*-test, *t*(12) = −1.713, *p* = 0.121).

### Genome and transcriptome assembly and annotation

The *Portulaca amilis* v1 genome assembly was 403.89 Mb in length and extremely contiguous (L50/N50 = 5 scaffolds/42.598 Mb; L90/N90 = 9 scaffolds/39.183 Mb), with nine primary scaffolds representing the nine expected chromosomes of the haploid *P. amilis* genome (Fig. 2a) based on the karyotype of a close relative (*P. grandiflora*^53^). The scaffolds were 92.9% complete and single copy as measured by the BUSCO v5 embryophyta database (C:96.9% [S:92.9%, D:4.0%], F:1.5%, M:1.6%, n:1,614), and eukaryotic BUSCOs were similarly complete (C:98.0% [S:82.7%, D:15.3%], F:1.2%, M:0.8%, n:255). Slightly less than half of the assembly was masked as repetitive elements (46.96%), which primarily consisted of LTRs (41.96%), DNA elements (29.36%), and LINEs (13.85%) (Fig. 2b). A repeat landscape of these elements did not show any sudden bursts of element activity or multiple peaks (Fig. 2b), which can signify genomic upheaval from hybridization or polyploidy events^54, 55^. The final annotation contained 53,094 gene models with 58,732 unique coding sequences (Fig. 2c).

**Figure 2.**
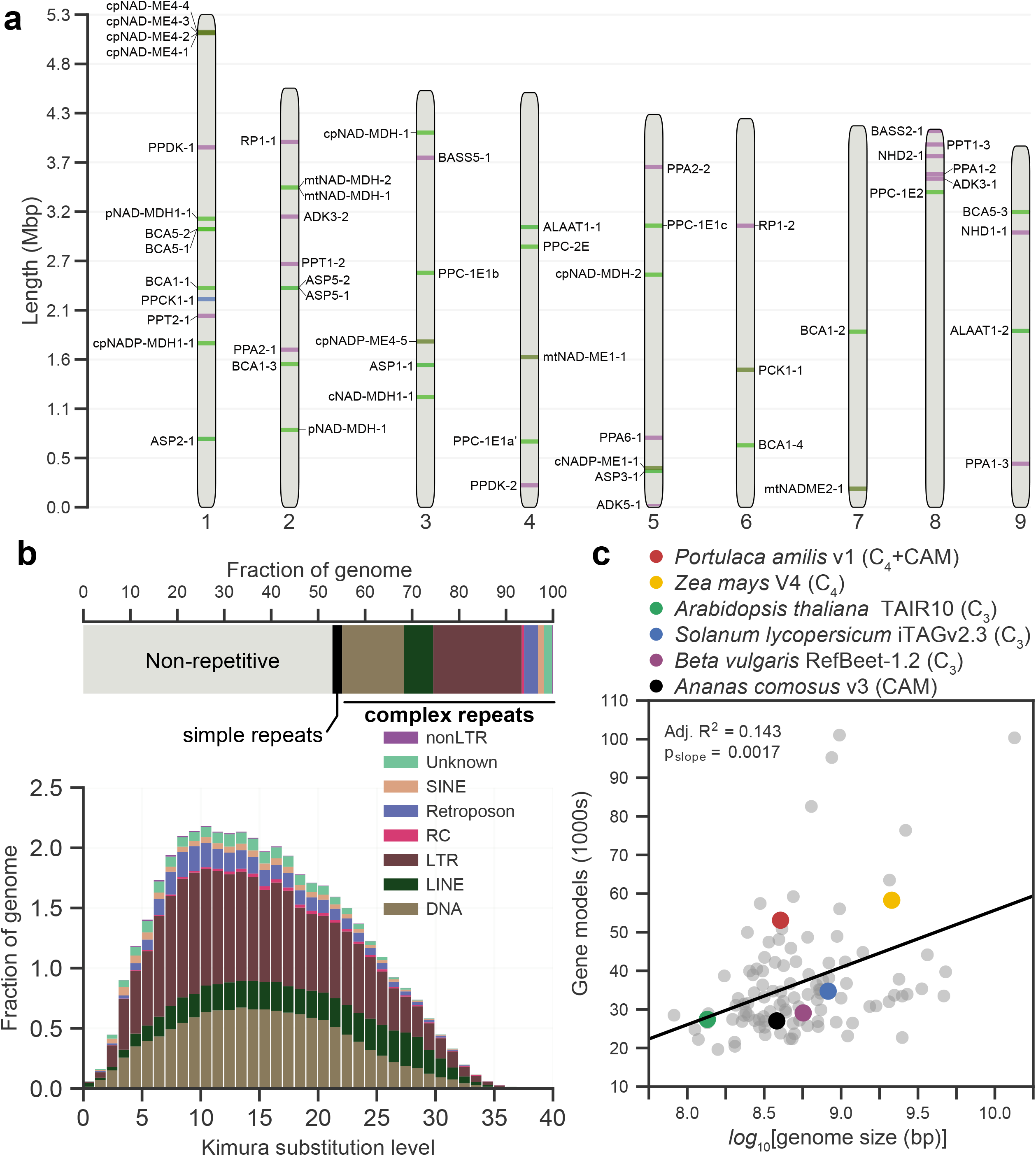
Idiogram of the nine primary scaffolds of the *P. amilis* genome with key C_4_ and CAM genes highlighted (a). Breakdown of repetitive elements of the *P. amilis* genome (b); the horizontal bar chart shows the fractions of major classes of repetitive elements and the repeat landscape (histogram) shows the relative abundances of repetitive elements versus the Kimura substitution level from the consensus. Scatterplot of number of gene models versus the log10-transformed genome size for 107 angiosperm genomes (c) with notable benchmarking and CCM genomes highlighted; data is from Zhao and Schranz^120^. Line in (c) shows results of ordinary least squares regression; one sample *t_slope_*(106) > 0, *p_slope_* = 0.0017.

The *P. oleracea de novo* transcriptome assembly contained 413,658 transcripts corresponding to 230,086 “unigenes” (hereafter, “genes”) that were highly complete but infrequently single copy: BUSCO v5 embryophyta (C:94.7% [S:16.4%, D:78.3%], F:3.2%, M:2.1%, n:1,614), eukaryota (C:98.9% [S:16.9%, D:82.0%], F:0.0%, M:1.1%, n:255). The high number of duplicate BUSCOs was expected, as most members of the *P. oleracea* species complex are polyploid^56^. Less than half of the transcripts (198,026) were predicted to be coding sequences, which corresponded to 45,285 genes.

A total of 21,621 *P. amilis* and 44,532 *P. oleracea* genes passed filtering for differential abundance analysis^57^, and over half of these genes (*P. amilis*, 62%; *P. oleracea*, 51.0%) were found to be significantly differentially abundant between well-watered and drought conditions following Benjamini Hochberg corrections^57^ for false discovery (*q* < 0.01; see Methods for details).

### Co-expression network analysis

The well-watered and drought *P. amilis* photosynthetic gene networks (PGNs) were similar in size and density, but had different distributions of node degree due to a small number of extremely highly connected genes belonging to modules paWW1 and paWW3 (Fig. 3a, Supplementary Fig. 1c, and Supplementary Table 1). The 104 genes of paWW1 represented all functional groups except starch synthesis, and paWW1 was characterized by C_4_-like expression: very high morning abundance that rapidly tapered off by late afternoon. Indeed, paWW1 contained at least one homolog of all core elements of the C_4_ pathway; including *paPPC-1E1a’*, the C_4_-specific *PPC* ortholog used by *P. oleracea*^28, 40, 41^ (Fig. 3c). The most similar drought module to paWW1 in terms of constituent genes, paD6, was 31% smaller (72 genes), while the largest drought module (paD1) was characterized by dark period peak expression and consisted primarily of light response and circadian transcription factors, and starch catabolism-related genes (Fig. 3b). Similar to *P. amilis*, the largest well-watered *P. oleracea* module (poWW2) contained *poPPC-1E1a’*; however, the *P. oleracea* transcriptome contained five *poPPC-1E1a’* transcripts distributed between two modules with C_4_-like expression (poWW2 and poWW1) (Fig. 3e). There was a greater size discrepancy between the *P. oleracea* PGNs, and the drought PGN was much larger and denser than the well-watered PGN (Fig. 3e, f, and Supplementary Table 1).

**Figure 3.**
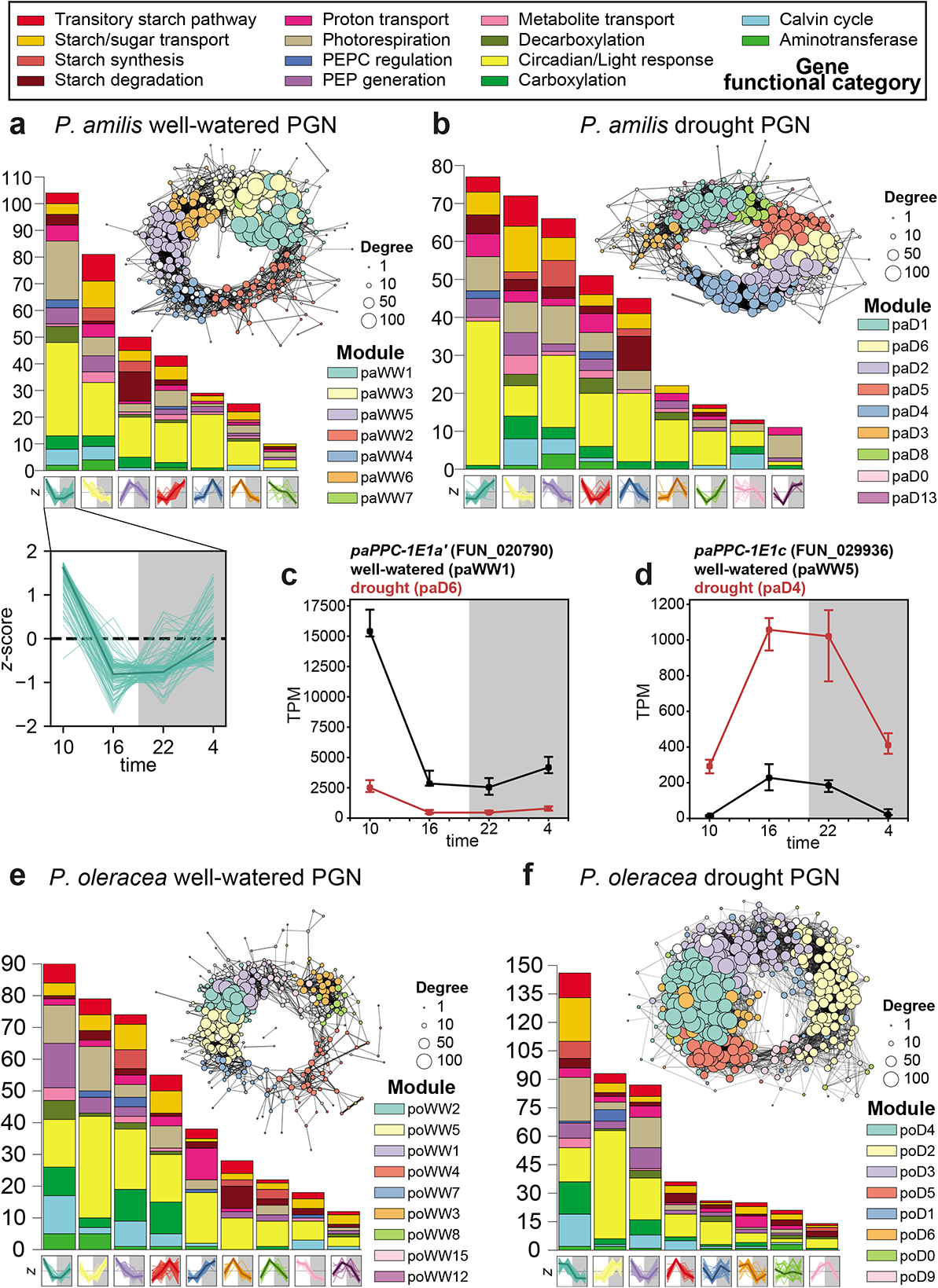
Photosynthetic gene networks (PGNs). Well-watered and drought PGNs of *P. amilis* (a-b), and *P. oleracea* (e-f) colored by WGCNA co-expression module identity. Each node in the graph represents one gene and node sizes represent their degrees. Each module’s size and functional composition is shown in the histogram, and the *z*-score normalized expression of each module’s constituent genes is shown along the horizontal axis. An example of the *z*-score normalized expression for module paWW1 is enlarged in (a) to show individual gene expression with the median highlighted in bold. Transcripts per million (TPM) normalized expression profiles of two focal *PPC* transcripts (*paPPC-1E1a’* and *paPPC-1E1c*) are shown in (c) and (d), respectively; points show median of size biological replicates and whiskers show interquartile range; black and red lines represent well-watered and drought samples, respectively.

We expected the CAM cycle to be broken into two co-expression networks: one that builds PEP from transitory starches and carboxylates CO_2_ into malate via BCA, PEPC, and MDH from dusk until dawn, and a second that decarboxylates malate into CO_2_ for fixation by RuBisCO during the day (Fig. 1b). As reported in *P. oleracea*^28, 40, 41^, *P. amilis* used the *PPC* paralog *paPPC-1E1c* for CAM (Fig. 3d), and this gene belonged to smaller modules with abundance profiles that peaked across the light-dark transition, and were dominated by starch degradation and circadian clock genes (paWW5 and paD4; Fig. 3a, b, d). Extracting the genes that remained co-expressed (preserved) across well-watered and drought conditions resulted in 28 genes of the *paPPC-1E1c* modules (42%) (Fig. 4a). With the exception of one photorespiratory gene (*paCAT1-2*, FUN_025647), all had functions related to carboxylation (*paPPC-1E1c* and *paBCA1-2*), starch metabolism and catabolism, or circadian rhythm and light response. At least one paralog of all members of the starch degradation pathway were preserved, except amylase (*AMY*) and beta-amylase (*BAM*), but one *BAM* homolog (*paBAM3-2*, FUN_041354) was recovered exclusively in the drought module, along with one paralog of aluminum activated malate transporter 4 (*paALMT4-2*, FUN_033599), which imports malate into the vacuole. However, no *MDH* homologs were recovered in this nocturnal CAM network, and most were recovered in drought modules paD6, paD2, and paD5 with C_4_-like expression.

**Figure 4.**
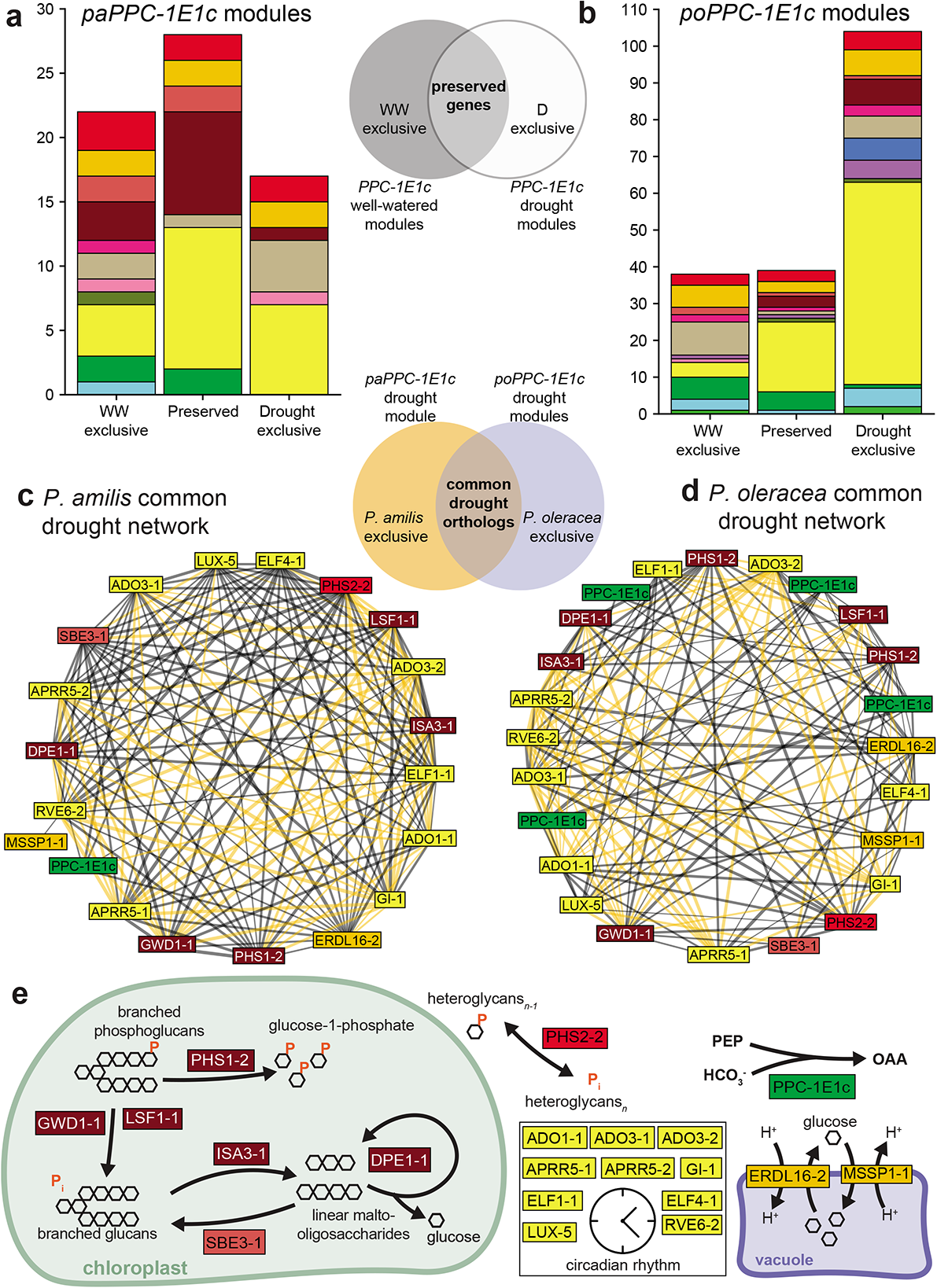
*PPC-1E1c* PGN preservation and subnetworks of common genes. Stacked bar charts showing the number and functional composition of genes that were exclusive to the well-watered or drought *PPC-1E1c* PGNs, or preserved across treatments in *P. amilis* (a) and *P. oleracea* (b). Subnetworks induced by genes common to both *P. amilis* (c) and *P. oleracea* (d) drought *PPC-1E1c* PGNs; drought PGNs include preserved and drought exclusive genes. Orange edges represent correlations among genes shared between species, while black edges are unique to each respective species. Metabolic pathways of the shared *PPC-1E1c* subnetworks (e), showing PEPC, sugar transport, and starch metabolism. Colors of bar plot units (a and b) and gene boxes (c-e) indicate functional categories as in Fig. 3.

The *P. oleracea* transcriptome contained multiple transcripts of *poPPC-1E1c* that were recovered in two well-watered modules (poWW4 and poWW8) and three drought modules (poD2, poD5, and poD9) that were consistent with use in CAM (Fig. 3e, f). The preserved genes were functionally similar to those preserved in the *P. amilis* nocturnal CAM network, and were involved with starch metabolism and catabolism, and the circadian clock (Fig. 4b). Among the genes preserved with *poPPC-1E1c* was cytoplasmic *poNAD-MDH1* (TRINITY_DN7462_c3_g1), which was the most highly expressed *NAD-MDH* under drought. Furthermore, many of the genes that link phosphorylative starch degradation to PEP generation were preserved in the nocturnal CAM network, including *AMY*, fructose-bisphosphate aldolase (*FBA*), glyceraldehyde-3-phosphate dehydrogenase (*GAP*), and phosphofructokinase (*PFK*). Although not preserved, the *poPPC-1E1c* drought modules included copies of phosphoglucomutase (*PGM*) and enolase (*ENO*), the final two steps in PEP generation from starch or soluble sugars.

We used the *paPPC-1E1c* and *poPPC-1E1c* drought modules to identify the set of genes common to both nocturnal CAM networks (Fig. 4c, d). By taking the intersection of the two sets of *PPC-1E1c* drought module genes (i.e., preserved and drought exclusive genes), we uncovered 21 *P. amilis* and 24 *P. oleracea* orthologs (some genes had multiple paralogs in the *P. oleracea* network). These orthologous nocturnal CAM networks contained one gene involved in starch synthesis (starch branching enzyme 3, *SBE3*), five phosphorylative starch degradation genes, and alpha-glucan phosphorylase 2 (*PHS2*), which is also part of phosphorylative starch metabolism and catabolism (Fig. 4e). These orthologous nocturnal CAM networks covered much of the phosphorylative starch degradation pathway that forms the basis of PEP generation in most CAM plants^58, 59^, but the interactions among many genes were different between species (Fig. 4c, d). The largest group of shared genes were core circadian clock elements: *Adagio* (*ADO*), also known as *ZEITLUPE* (*ZTL*), *1* and *3*, two *Arabidopsis pseudo-response regulator 5* (*APRR5*) paralogs, *EARLY FLOWERING* (*ELF*) *1* and *4*, *GIGANTEA* (*GI*), one *LUX ARRHYTHMO* (*LUX*) paralog, and one *REVEILLE 6* (*RVE6*) paralog (Fig 4c, d, e). Although the two species’ nocturnal CAM networks shared many key orthologs, there were notable differences. The *P. oleracea* network did not contain an *ALMT* homolog, but did contain three paralogs of *PEPC kinase 1* (*PPCK1*) that were all highly abundant and significantly upregulated under drought (*q* < 1.0 x 10^-7^) (Supplementary Fig. 4). PPCK increases the activity of PEPC, which is inhibited by malate, forming a negative feedback loop^60^. While all three *poPPCK* transcripts were upregulated under drought with peak nighttime expression indicative of use in CAM, all had higher daytime levels of abundance under drought as well—which points towards a dual role in both C_4_ and CAM. The single *PPCK* ortholog in *P. amilis* (FUN_002684) was also significantly more abundant under drought (*q* = 0.0011) and had peak nocturnal expression (Supplementary Fig. 3), but it was recovered in drought module paD1, which increased in abundance throughout the night (Fig. 3b). PPCK activity is typically essential for highly active CCMs, and a CAM-like expression pattern in *paPPCK1* would suggest low PEPC activity during C_4_. However, we found a derived amino acid residue, glutamic acid (E), in *paPPC-1E1a’* at the same location where an aspartic acid (D) residue (D509) was demonstrated to reduce malate inhibition in *Kalanchoë* and *Phalaenopsis* orchids^61^. In *Kalanchoë* and *Phalaenopsis*, the D509 residue was shown to be derived from either arginine (R), lysine (K), or histidine (H), the latter of which is present at position 509 in *poPPC-1E1a’*. Glutamic acid is functionally similar to aspartic acid, and may similarly reduce malate inhibition; thus, allowing for high PEPC activity during C_4_ without the need for PPCK.

We considered two modules each in *P. amilis* (paWW1 + paWW3 and paD5 + paD6) and *P. oleracea* (poWW2 + poWW1 and poD3 + poD4) when calculating the preservation of the C_4_ pathway because these pairs of modules had C_4_-like expression patterns and were strongly connected (Fig. 3a, e). In the *P. amilis* C_4_ PGNs, 86 genes (38%) were preserved (Supplementary Fig. 4), which represented of all core C_4_ (*ALAAT*, *ASP*, *BCA*, *ME*, *MDH*, *PPC-1E1a’*) and PEP regeneration gene families (adenylate kinase (*ADK*), sodium/pyruvate cotransporter (*BASS*), sodium-hydrogen antiporter (*NDH*), soluble inorganic pyrophosphatase (*PPA*), pyruvate, phosphate dikinase (*PPDK*), and PEP/phosphate translocator (*PPT*)). It also contained members of the entire transitory starch pathway that builds and transports various sugars and carbohydrates (e.g., *ENO*, *GAP*, phosphoglycerate kinase (*PGK*), *PGM*, and triose phosphate/phosphate translocator (*TPT*)). In addition to *paBASS2-1* (FUN_026309), which imports pyruvate into the chloroplast as a substrate for PEP generation, we recovered two paralogs of methyl erythritol phosphate (*MEP*), also known as *RETICULATA-RELATED* (*RER*), *4* (*A. thaliana* ortholog AT5G12470). MEP transporters have been implicated as mesophyll pyruvate importers and bundle sheath pyruvate exporters in maize^62, 63^, and thus may serve a similar function to BASS. The preserved modules also contained key enzymatic regulators (e.g., serine/threonine-protein phosphatase (*PP2A*) subunits and RuBisCO activase (*RCA*)) and transcription factors and clock elements (e.g., *CIRCADIAN CLOCK ASSOCIATED 1* (*CCA1*)).

*Portulaca amilis* and *P. oleracea* span a fairly deep node in the *Portulaca* phylogeny (Fig. 1c) and it was unclear which facets of C_4_ they might share. A similar fraction of C_4_ module genes were preserved between well-watered and drought C_4_ PGNs in *P. oleracea* (117; 41.2%), and these largely consisted of the same gene families (58.3% overlap with preserved *P. amilis* C_4_ gene families). To identify the orthologous components of the *P. amilis* and *P. oleracea* C_4_ gene networks, we took the intersection of their well-watered C_4_ PGNs, which contained 186 and 165 homologs, respectively, that represented 258 unique ortholog groups. Roughly one quarter (69/258; 26.7%) of these ortholog groups were recovered in the common C_4_ networks, while 117 (45.3%) and 72 (27.9%) were exclusive to *P. amilis* and *P. oleracea*, respectively. The common networks included many of the most highly expressed copies of key C_4_ genes, including the PEP generation pathway, and most of the photorespiratory pathway (Supplementary Fig. 5).

Indicative of the unique evolutionary histories of *P. amilis* and *P. oleracea*, each species’ C_4_ PGN contained exclusive orthologs of most C_4_ gene families, some of which represent species-specific duplication events. Both species exhibited exclusive use of orthologs of *ADK*, *ALAAT*, *ASP*, *BCA*, *MDH*, *ME*, *NDH*, *PP2A*, *PPA*, mitochondrial uncoupling protein (*PUMP*), RuBisCO, and *RCA*, among others. Some exclusive homologs fit biochemical expectations. Exclusive to *P. amilis* (NADP-type C_4_) were three chloroplastic *NADP-ME4* paralogs (FUN_006401, FUN_006404, and FUN_006411) which did not have orthologs in the *P. oleracea* transcriptome. These three copies were on the same strand and physically close in the genome (within ∼85kb); they possibly represent tandem duplication events of *paNADP-ME4-4* (FUN_006423), which is ∼50kb downstream of FUN_006411 (Fig. 2a). In *P. oleracea*, two *ASP* (TRINITY_DN71325_c0_g1 and TRINITY_DN346_c1_g3) and three *ALAAT* (TRINITY_DN43865_c0_g1, TRINITY_DN43865_c0_g2, and TRINITY_DN81553_c0_g1) homologs were found in orthogroups that had no *P. amilis* members. Many of the genes that were exclusive to either C_4_ PGN were circadian clock and light response transcription factors, or regulatory enzymes. For example, unique orthologs of the *FAR-RED IMPAIRED RESPONSE* family were part of each C_4_ PGN, while only the *P. amilis* C_4_ PGN contained an *APRR* homolog, and only the *P. oleracea* C_4_ PGN contained a *LATE ELONGATED HYPOCOTYL* (*LHY*) transcription factor.

We used the drought C_4_ PGNs to identify orthologous ancestral elements of the daytime portion of the CAM cycle. We expected this portion of the CAM cycle to be nearly indistinguishable from the C_4_ cycle (Fig. 1a, b), but to only exhibit C_4_-like expression under drought stress. To be considered part of the ancestral CAM cycle, elements needed to both be shared between the two drought C_4_ PGNs and be significantly upregulated under drought (*q* < 0.05). Eleven orthologs met these potentially CAM-induced criteria, but only one had a function directly related to daytime CO_2_ release from nocturnal malate stores: chloroplastic *NADP-ME4-5* (FUN_015913; TRINITY_DN81025_c0_g1). *PUMP5-1*—also known as dicarboxylate transporter 1 (*DIC1*)—of both species also appeared CAM-induced, but was not significantly differentially abundant when considering the entire time series in *P. oleracea* (*q* = 0.24). The *P. amilis* C_4_ PGNs contained a drought exclusive mitochondrial *NAD-ME2-1* (FUN_037145) that was significantly upregulated (*q* < 0.01), and a second, preserved ortholog (*NAD-ME1-1*; FUN_021996) that was significantly upregulated under drought (*q* < 0.05). Coupled with upregulated *PUMP5-1* expression, this suggests that the NAD decarboxylation pathway, which occurs mainly in the mitochondria, may still be elicited by CAM in *P. amilis* despite NADP-type C_4_. In contrast, constitutive expression of the same mitochondrial orthologs in *P. oleracea* (*poNAD-ME1-1* and *poNAD-ME2-1*) is consistent with use in C_4_, suggesting that *P. oleracea* may have recruited its ancestral CAM NAD-decarboxylating pathway into its C_4_ system. Finally, and unexpectedly, one *P. amilis* paralog of the RuBisCO small subunit B (*paRBCS-B-1*; FUN_016561) fit a CAM induced abundance pattern (*q* < 0.001) and was expressed on the same order of magnitude as other *RBCS* homologs.

### Motif identification and enrichment

Enriched motifs were detected in at least one region (i.e., 3’UTR, 5’UTR, introns, or upstream promoter) for gene models in many *P. amilis* modules. However, no enriched motifs were identified for paWW1 or paWW3—which contained the bulk of C_4_ genes when well-watered. In contrast, the 5’UTRs and upstream promoter regions of the preserved *paPPC-1E1c* module were enriched in 10 unique motifs (*p* < 0.01), all of which had circadian or light response functions (Table 1). These motifs were all annotated to Myb-related families, except bHLH74, which contains a G-box motif (CACGTG), and governs flowering time in response to blue light^64^. All enriched motifs contained either an Evening Element (EE; AAATATCT) or GATA (HGATAR) motif. EE and GATA motifs often serve as binding sites for core circadian oscillator transcription factors such as RVEs, CCA1, LHY, and APRR1, also known as TIMING OF CAB 1 (TOC1)^65–67^.

**Table 1.**
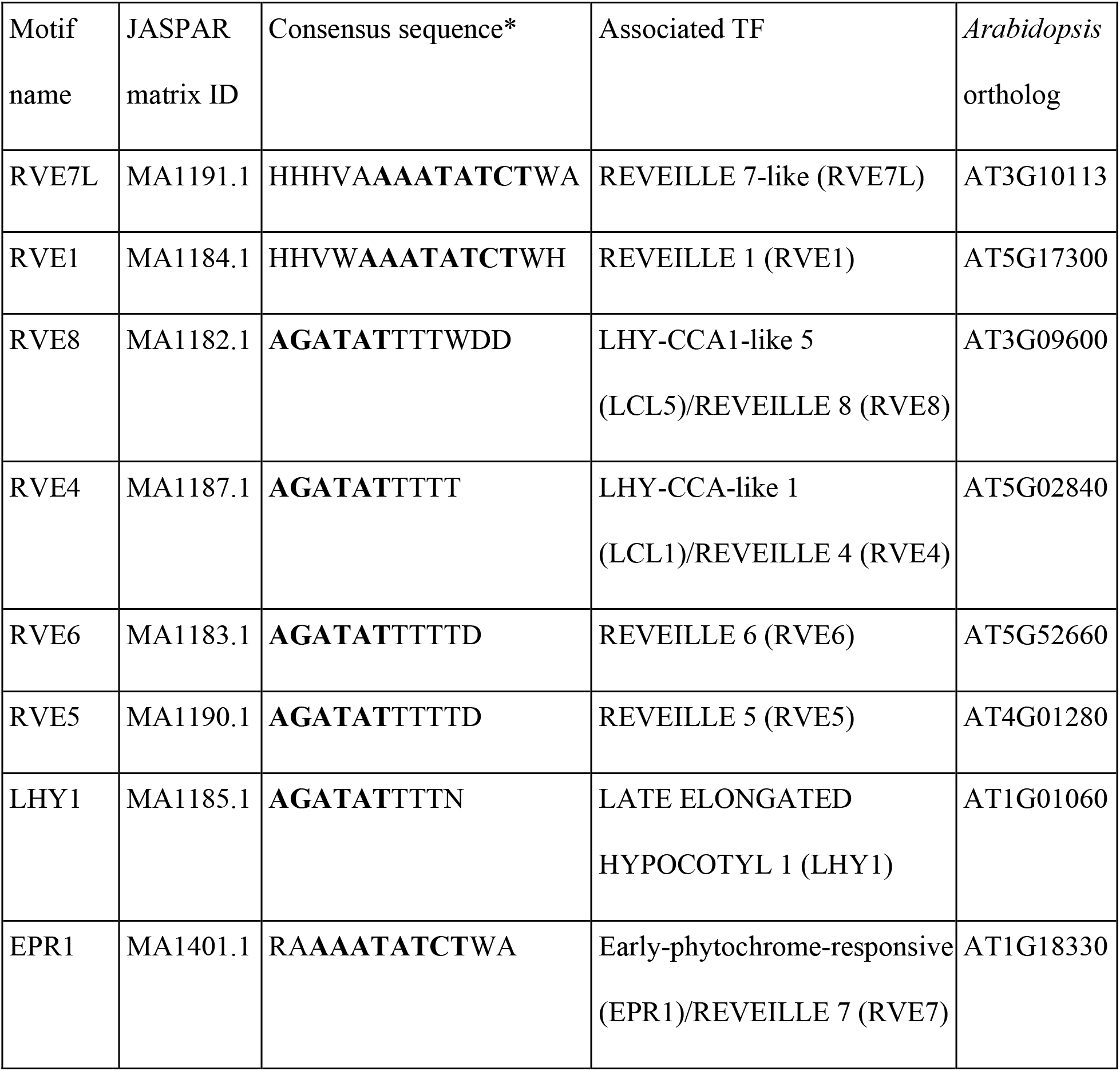
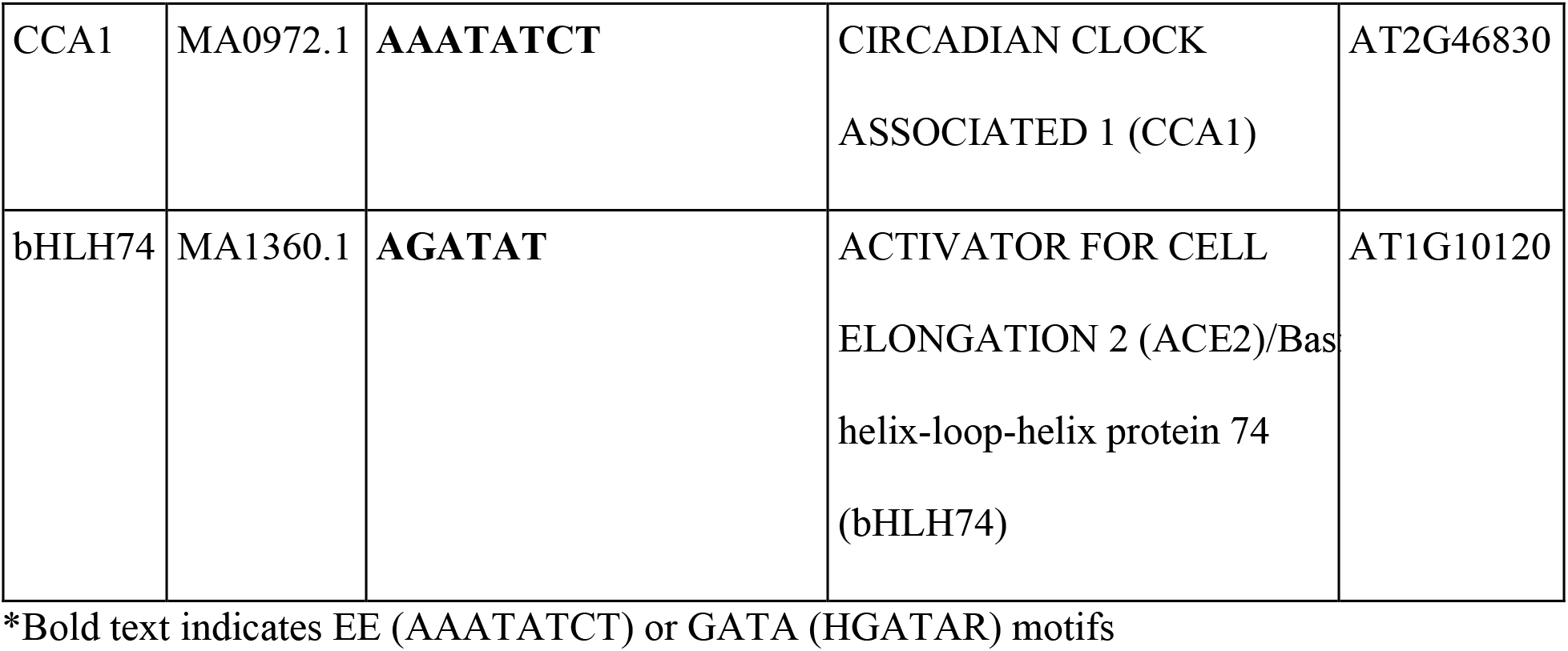
Motifs enriched in the 5’UTR of genes preserved in the *paPPC-1E1c* module

There were no enriched motifs in or near genes preserved with *paPPC-1E1a’*, and the *P. amilis* C_4_ PGN genes were only enriched in four motifs: one Myb (MA1028.1), two bZIP (MA0968.1, MA1033.1), and one AP2/ERF (MA1261.1) factor (Supplementary Table 2). While these motifs were significantly enriched (*p <* 0.01), they were much less widespread than the enriched motifs found in the preserved CAM genes. At the low end, the KAN4 motif (MA1033.1) was present in just five 5’UTRs (8.5% of sequences), and the most widespread (bZIP68, MA0968.1) was present in 12 sequences (20.3%). Motifs found in the preserved *paPPC-1E1c* genes were present in 20.7-58.6% of the sequences, and all 10 enriched motifs were found at least once in the 5’UTR and promoter region of *paPPC-1E1c*.

## Discussion

Plant photosynthetic adaptations can serve as models for exploring central themes of biology, such as the evolution of modularity and evolvability, gene and genome duplication, convergent evolution, and the evolution of novel phenotypes. Network analyses provide powerful tools for identifying and comparing modules of genes related to functions of interest against a background of tens of thousands of genes with complicated temporal expression patterns. C_4_+CAM photosynthesis has only been reported in three plant lineages, *Portulaca*^29^, *Spinifex*^30^, and *Trianthema*^31^, but CCMs have evolved many dozens of times^8–12^ and the use of multiple CCMs has been documented in aquatic plant lineages^68, 69^. CCM evolution is a classic example of the repeated assembly of a new function via the integration of multiple, existing gene networks^7^. When coupled with patterns of transcript abundance and placed in an evolutionary context, our network results provide an initial interpretation of how these metabolic pathways interact in *Portulaca,* and how they were evolutionarily assembled.

### The evolution of C_4_+CAM in Portulaca

Facultative CAM is ancestral in *Portulaca*, and likely evolved by coupling elements of the TCA cycle (i.e., PEPC and BCA) to the phosphorylative starch degradation pathway via the circadian clock. CAM in both species was restricted to a minimal set of tightly regulated nocturnal pathways that linked *PPC-1E1c* to the transitory starch pathway, with only parts of the decarboxylation pathways distinguishable from C_4_. This aligns with previous hypotheses that the initial steps in CAM evolution do not require “rewiring” of gene networks, but rather occur by upregulating nocturnal organic acid production that occurs in all plants^13^. Subsequent canalization of these fluxes occurs by bringing these networks under circadian control. If C_4_ evolution began in *Portulaca* while the nighttime portion of CAM included derived gene networks, but the daytime portion did not, CAM would have likely imposed few pleiotropic constraints given redundant paralogs, because recruitment of genes into C_4_ would not have required modification of existing CAM decarboxylation pathways in the mesophyll. Sufficient paralogs of NAD- and NADP-type enzymes were present in the most recent common ancestor of *P. amilis* and *P. oleracea* to allow C_4_ and CAM to use separate genes, and for differential enzyme co-option during their independent C_4_ origins. However, transcriptomic data presented here and elsewhere^40, 44^ imply that CAM-specific decarboxylation pathways did exist in the common ancestor to *Portulaca*. It therefore seems likely that the initial steps towards C_4_ evolved within a particular window of CAM evolution before other aspects of CAM were fixed, such as diurnal stomatal closure or changes to leaf ultrastructure that greatly reduced intercellular diffusion of metabolites.

Shared use of some enzymes and pathways suggests that the most recent common ancestor of *Portulaca* may have had some C_4_ characters. Shared use of photorespiratory modules, found here, and proto-C_4_ anatomy in C_3_-C_4_ intermediate taxa^49, 50^ suggest that C_2_ metabolism—the enhanced refixation of photorespired CO_2_ by preferential expression of various photorespiratory enzymes^25, 70^—may also be ancestral to *Portulaca*. Like other CCMs, C_2_ requires the recruitment of existing gene networks, and our comparison of well-watered C_4_ gene networks revealed many shared photorespiratory orthologs that may point to a single origin. However, it is difficult to distinguish between a scenario in which these orthologs were co-opted into C_2_ metabolism once near the stem of extant *Portulaca*, or more recently along multiple branches in parallel. Shared use of PEPC-1E1a’ in C_4_ has been hypothesized to have occurred through parallel recruitment^28^, stemming from “preconditioning”, such as increased expression^71^ or adaptive amino acid substitutions^29^. C_2_ metabolism is hypothesized to typically be followed by rapid evolution of a C_4_ system^25, 70^, and therefore represents a limit of what we expect to be shared among *Portulaca* lineages with distinct morphologies and biochemistries. Broader sampling across the *Portulaca* phylogeny is needed to precisely delimit the shared initial steps towards C_4_ and subsequent parallel refinement of C_4_+CAM.

### Integration of C_4_ and CAM in Portulaca

*Portulaca amilis* and *P. oleracea* have canonical C_4_ pathways, but constitutively operate very weak CAM cycles (at least at the transcriptional level) that are upregulated under drought. The core elements of C_4_ and CAM were moderately preserved in both species, and where parts of the C_4_ and CAM pathways could theoretically overlap, we often found alternative homologs. Increased CAM activity was associated with a reduction in size of C_4_ PGNs, but there was no evidence that this was due to dual roles of genes in nocturnal CAM physiology under drought. If transcript abundances of C_4_ and CAM related genes can serve as a proxy for enzymatic and metabolic activity, conflicting intracellular physiological cues from CAM likely played a minor role in altering C_4_ activity. Given the strong diffusion gradients between mesophyll and bundle sheath of C_4_ species^72^, we suspect that the evolution of C_4_ in *Portulaca* has resulted in two-cell CAM, as has been previously hypothesized^37^, where malate produced by CAM in the mesophyll is largely decarboxylated in the bundle sheath, but it is unclear if overnight storage occurs in the mesophyll, bundle sheath, or both (Fig. 5). Drought-induced upregulation of decarboxylating enzymes was consistent with expectations given C_4_ biochemistries. Upregulation of *paNAD-ME1-1* and *paNAD-ME2-1* during CAM induction in *P. amilis* indicated that C_4_ did not co-opt the entirety of the diurnal portion of the CAM cycle (Fig. 5). In contrast, constitutive use of *poNAD-ME1-1* and *poNAD-ME2-1* in *P. oleracea* was consistent with recruitment and integration of the ancestral CAM NAD-decarboxylating pathway into C_4_, while the NADP-pathway maintained its role in CAM (Supplementary Fig. 6).

**Figure 5.**
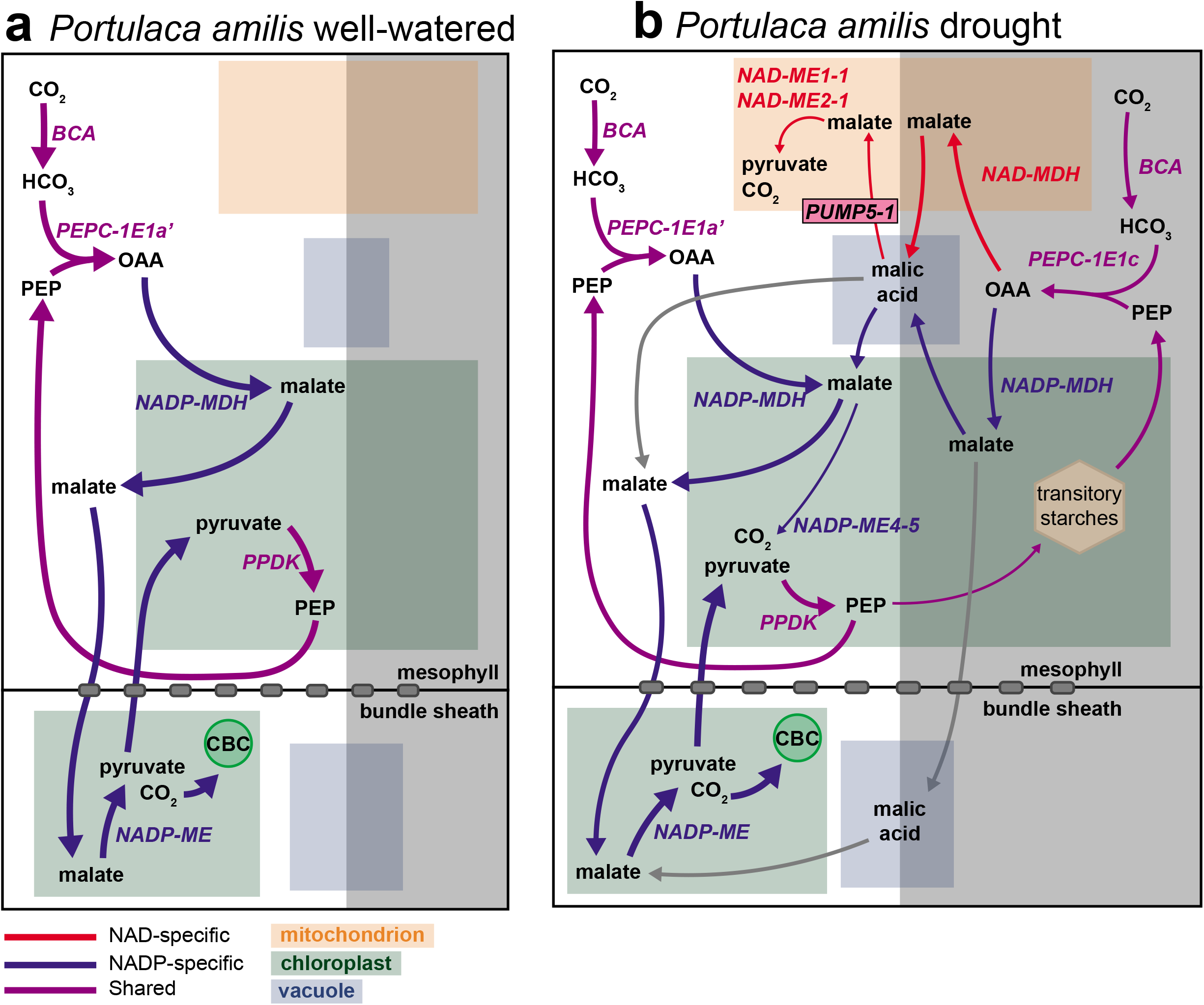
Hypothesized biochemical cycles of C_4_+CAM in *P. amilis* when well-watered (a) and droughted (b). Red, blue, and purple lines indicate NAD-specific, NADP-specific, and shared reactions, respectively, and grey lines show novel, possible pathways for malate that are unique to C_4_+CAM photosynthesis. Line thicknesses are indicative of relative fluxes through pathways. ALAAT, alanine aminotransferase; ASP, aspartate aminotransferase; BCA, beta carbonic anhydrase; CBC, Calvin-Benson cycle; NAD-MDH, NAD-dependent malate dehydrogenase; NADP-MDH, NADP-dependent malate dehydrogenase; NAD-ME, NAD-dependent malic enzyme; NADP-ME, NADP-dependent malic enzyme; PEP, phosphoenolpyruvate; PEPC, PEP carboxylase; PPDK, pyruvate, phosphate dikinase; OAA, oxaloacetate.

Our network results confirmed that the nocturnal portion of CAM is a fixed gene network—rather than the result of drought-induced rewiring—that is both transcriptionally and post-transcriptionally regulated under drought to increase CAM activity. Gas exchange measurements in *Portulaca* species show substantial net CO_2_ loss during the night when well-watered^39–42^. Therefore, well-watered expression levels of the preserved CAM-related genes are either not high enough to re-fix all nocturnally respired CO_2_ or must have their products regulated at a post-transcriptional level. Evidence for the latter is given by expression patterns of PPCK, which increases the activity of PEPC^60^. *PPCK* was upregulated in both species under drought, and during the dark period in particular, which implies a role in increasing nocturnal PEPC activity for CAM under drought. In *P. oleracea*, the maintenance of high daytime *poPPCK* expression with a nocturnal peak in abundance suggests that PPCK may play a dual role in both C_4_ and CAM. In contrast, we were surprised to find that *paPPCK* had expression only consistent with use in CAM, as we are not aware of other C_4_ species without high morning expression of *PPCK*; but a derived amino acid residue in PEPC-1E1a’ has the potential to increase its activity by reducing malate inhibition, as has been show in *Kalanchoë* and *Phalaenopsis*^61^. Therefore, the avoidance of pleiotropic effects has occurred through redundant homologs (e.g., decarboxylating enzymes) and adaptive mutations, but *P. oleracea* provides evidence that pleiotropy may not constrain some genes.

### Portulaca as a model for metabolic evolution

The natural history of *Portulaca* provides a compelling model system for studying many evolutionary and ecophysiological questions, and its unique metabolism and experimental practicality make it a powerful tool for genetic engineering. Forward engineering of both C_4_^73^ and CAM^74^ have become priorities for agriculture as arid lands expand and populations grow. *Portulaca* comprises over 100 species that are found in nearly every country and have invaded some continents multiple times^56^. Despite the small genome size of *P. amilis*, duplication events deeper in the history of the Portulacineae^75^ have played large roles in the evolution of C_4_+CAM, with implications for neofunctionalization and the evolution of modularity. With the generation of the first *Portulaca* genome, we provide a resource for studying these past events and for the functional genetic work to understand and engineer CCMs.

CAM is found in every major clade of the Portulacineae to varying degrees^34, 43, 45, 46, 76, 77^ (Fig. 1c), and stronger and constitutive CAM have evolved in *Portulaca*’s closest relatives, Anacampserotaceae and Cactaceae^43, 78, 79^. These three clades overlap in climate space^77^ and in many parts of their geographic ranges^80^, exhibit succulent morphologies capable of photosynthesis in their stems^43, 81^, and likely share the same ancestral CAM cycle. It remains an open question why only *Portulaca* evolved C_4_+CAM, while stronger CAM evolved in close relatives. C_4_ and CAM consume more energy than C_3_ photosynthesis, and we lack models to predict environmental conditions that would select for C_4_ in a CAM plant (or vice versa). Selective pressures to evolve C_2_ in the facultative CAM ancestor of *Portulaca* may have been no different than in C_3_ plants, as CAM does not necessarily reduce photorespiration, and in some cases can exacerbate it^20^. Future progress on these topics will be made through continued study of the C_3_-C_4_ Cryptopetala clade that uses C_2_ metabolism.

Despite an increased general interest in CCMs over recent years, and *Portulaca*’s C_4_+CAM system in particular, we have only a limited understanding of the benefits of combining C_4_ and CAM. The advantages of facultative CAM vary between lineages, as does carbon gain, which is typically <10% of the daytime photosynthetic activity^82, 83^. Facultative CAM increases water use efficiency and can prevent reduced growth and reproductive output under drought while also reducing photoinhibition (radiation induced damage to photosystems)^82, 83^. Measures of chlorophyll *a* in *P. oleracea* over a water limitation experiment provide evidence that facultative CAM reduces photoinhibition^40^. A photoprotective role of CAM is further corroborated by rapid recovery of photosynthetic rate upon rewatering^38, 39^, indicating little or no damage is done to photosystems. Gas exchange measurements also demonstrate that *Portulaca* continue to grow despite several days of drought stress^38, 39^. Furthermore, drought stress may increase CO_2_ leakiness of bundle sheath in C_4_ plants, a problem augmented when carboxylation activity decreases relative to decarboxylation^84^. Facultative CAM may provide some carbon and photoprotection during drought that accelerates the transition to full photosynthetic capacity upon rewatering and allows for growth and reproduction despite large water deficits.

The annotated *P. amilis* genome and metabolic modules identified here, combined with recent resources for transgenic research in *Portulaca*^85^ signal a significant shift toward the development of an experimental, functional genomics-based research program in C_4_+CAM. On one level, we suggest that *P. amilis* is a better candidate than *P. oleracea* for further model system development, in part because *P. oleracea* exists within the context of a larger species complex of over a dozen described species and subspecies with uncertain relationships and multiple whole genome duplication events^56^. *P. amilis* has an indistinguishable life history from *P. oleracea* (i.e., 3-5 week life cycle, thousands of seeds produced, self-fertilization), but does not suffer from the same taxonomic challenges and variations in ploidy. On the other hand, we view the *Portulaca* clade as providing not one, but at least three, evolutionary origins of C_4_+CAM systems, and there is much to be learned in understanding their similarities and differences.

## Conclusions

The evolution of modularity is a common theme in studies of evolutionary development and the origination of novel phenotypes. Network-based analyses can distinguish complex and highly overlapping metabolic modules, and establish their orthology between species. C_4_+CAM provides a unique opportunity to understand how a single module that is typically recruited into two distinct functions is regulated when both of the new functions operate in a single organism. Remarkably, their integration appears possible in part via additional modularization, with little overlap in C_4_ and CAM gene networks. Assignment of homologs to functions and generation of a high-quality genome for *P. amilis* were necessary first steps for future functional genetic work on C_4_+CAM. Perhaps the most pertinent questions surround the utility of C_4_+CAM, which could be addressed in part by knock-down or knock-out *PPC-1E1c* lines to study the fitness of a purely C_4_ *Portulaca*. In addition to functional genetic experiments, similar CAM-induction experiments are needed throughout *Portulaca* to establish homology between metabolic pathways across the multiple origins; especially in the C_3_-C_4_ Cryptopetala clade, which will provide greater clarity on which elements of C_4_+CAM are ancestrally shared across multiple origins in *Portulaca*.

## Materials and Methods

### Plant materials and CAM-induction experiment

Specimens of *Portulaca amilis* and *P. oleracea* were collected in Casselberry, Florida, U.S.A. and propagated in the Plant Environmental Center at Brown University. All studied *Portulaca* are selfing, and many are primarily cleistogamous^86, 87^; cleistogamous flowers self-pollinate in the developing flower bud without the need of a vector, and most buds progress directly to fruit set without opening. S2 seeds were germinated at Yale’s Marsh Botanical Garden. One S2 *P. amilis* was removed from soil, gently washed to remove soil, flash frozen whole in liquid N_2_, and shipped overnight to Dovetail Genomics (Scotts Valley, CA, U.S.A.) for whole genome sequencing and assembly. An S4 individual was collected and vouchered at the Yale University Herbarium in the Peabody Museum of Natural History.

S3 *P. amilis* and S2 *P. oleracea* were germinated in a growth chamber with the following environmental conditions: 00:00-06:00, dark, 22°C; 06:00-20:00, lights on, 25°C; 20:00-23:59, dark, 22°C. Plants were regularly bottom watered and the mean photosynthetically active radiation (PAR) at plant level was ∼385 mol · m^-2^ · s^-1^ during the 14 hour photoperiod. After plants had reached maturity and branched sufficiently (∼4 weeks), eight plants of each species were selected for CAM-induction: six biological replicates and two indicator plants. Indicator plants were monitored daily for fluctuations in titratable acidity to determine CAM induction, and were not included in any further analyses. To induce CAM, water was withheld from plants, and significant (one sample *t*-test, *p* < 0.01) increases in nocturnal titratable acidity were observed in *P. amilis* and *P. oleracea* indicator plants after seven and eight days, respectively. Leaf tissue was collected from all six biological replicates for titratable acidity assays at 16:00 and 04:00 (the following day) and bulk RNA at 10:00, 16:00, 22:00, and 04:00 (the following day) on day 0 (referred to as “well-watered”) and days 8 and 9 (referred to “drought”).

Titratable acidity was assessed by boiling 0.2-0.3g fresh leaf tissue in 60mL 20% EtOH until the volume was reduced by half (∼30mL). Distilled water was then added to return the volume to 60mL. This process was repeated, and the supernatant was then covered and allowed to cool to room temperature. Samples were then titrated to a pH of 7.0 using 0.002M NaOH, and recorded in units of μEq H^+^ per gram fresh mass—calculated as volume titrant (μL) ⨉ titrant molarity (M) / tissue mass (g).

Total leaf RNA was extracted using Option 2 (CTAB/TRIzol/sarkosyl) from Jordon-Thaden et al.^88^. Sample purity was measured using a NanoDrop (Invitrogen by Thermo Fisher Scientific, Carlsbad, CA), and a representative set of 12 samples were further assayed on using a 2100 Bioanalyzer (Agilent Technologies, Santa Clara, CA) for fragment length distribution and RNA integrity (RIN); RIN scores ≥ 7 were considered high quality. Library preparation included poly(A) tailing to pull down mRNA, and sequencing was done on an Illumina HiSeq (2×100bp paired-end reads), with an expected 25 million reads per sample. Four of the 96 total libraries failed during sequencing; all were *P. oleracea* samples (one 10:00 drought, one 16:00 drought, and two 04:00 drought).

### Genome and transcriptome assembly and annotation

The *Portulaca amilis* v1 genome was assembled *de novo* by Dovetail Genomics, using one Chicago^89^ and two Dovetail Hi-C^90^ libraries. The Chicago library was sequenced on an Illumina HiSeqX, resulting in 178 million 2×150bp paired-end reads that yielded 169.1x physical coverage of the genome (1-100kb pairs). The two Dovetail Hi-C libraries were also sequenced on an Illumina HiSeqX; libraries one and two contained 202 million and 342 million 2×150bp paired-end reads, respectively, which together provided 52,626.57x physical coverage of the genome (10-10000kb pairs). Meraculous 2^91^ was used to generate an assembly from 791 million paired-end shotgun read pairs (mean insert size ∼550bp and ∼350bp) with kmer size 91. The Meraculous assembly and shotgun, Chicago, and Hi-C reads were then used for iterative scaffolding using HiRise v2.1.6-072ca03871cc^89, 90^ to create the final assembly consisting of 4,053 scaffolds.

The *P. amilis* v1 genome was annotated using funannotate v1.6.0 (http://doi.org/10.5281/zenodo.3354704), a modular software package for genomic scaffold cleaning, genomic element prediction (i.e., genes, UTRs, tRNAs, etc.), functional annotation, and comparative genomics. All annotation steps after genome assembly were run within funannotate unless otherwise noted, and a step-by-step walkthrough of the entire annotation process can be found on GitHub (https://github.com/isgilman/Portulaca-amilis-genome).

Using funannotate’s ‘clean’ function, scaffolds were checked for duplicates with a “leave one out” strategy where, starting with the shortest scaffold, one scaffold was left out and mapped back to the remaining scaffolds with minimap2^92^. Scaffolds were sorted by size from largest to smallest using funannotate’s ‘sort’ function. Outside of funannotate, repetitive element analysis was conducted with RepeatModeler v1.0.11 and RepeatMasker v4.0.9-p2 (http://www.repeatmasker.org). RepeatModeler was used to create a custom repeat database for *P. amilis* and further classification of unknown repeats was done with TEclass v2.1.3^93^. The custom repeats were then passed to RepeatMasker, where complex repeats were soft masked. An additional round of masking was done using the Viridiplantae repeat database from RepBase (RepBaseRepeatMaskerEdition - 26 October 2018)^94^ and transposable element data from Dfam v3.0^95^. All repetitive elements were then cleaned using the RepeatMasker utility script ‘ProcessRepeats’.

Prior to *ab initio* gene prediction, we used funannotate’s ‘train’ function to create a preliminary set of high-quality gene models for training prediction software using the soft masked scaffolds and RNAseq reads from the CAM-induction experiment. Paired-end raw RNAseq reads were filtered by quality (phred < 25) and trimmed to remove adapters, poly-A/T tails longer than 12 bp, and low-quality tails (phred < 20) using PRINSEQ v0.20.4^96^. To capture the greatest diversity of transcripts and provide sufficient depth to call rare isoforms, we pooled all RNAseq reads from the six biological replicates across four time points and two conditions (well-watered and drought), totaling ∼1.58 billion reads from mature leaf tissue. RNAseq reads were then normalized using the ‘insilico_read_normalization.pl’ utility script within Trinity v2.8.5^97^. Funannotate’s ‘train’ function first builds a coordinate-sorted BAM file and associated index file using HISAT2 v2.1.0^98^ for an initial genome-guided transcriptome assembly with Trinity. The resulting initial transcripts then were fed through the PASA pipeline v2.3.3^99^ to generate gene models and predicted coding sequences using TransDecoder v5.5.0 (https://transdecoder.github.io). Kallisto v0.45.1^100^ was then used to quantify abundance data, and PASA gene models with abundance at least 10% of the most abundant gene model at each respective locus were retained for training the *ab initio* predictors Augustus v3.2.3^101^, SNAP v2013_11_29^102^, glimmerHMM v3.0.4^103^, and GeneMark-ET^104^.

Funannotate’s ‘predict’ function was then used to produce an initial set of gene models from the scaffolds using high abundance PASA models, *ab initio* prediction software, and evidence from transcripts and predicted proteins. First, transcript alignment information from the coordinate-sorted BAM file and protein alignments from Diamond v0.9.24^105^ and Exonerate v2.4.0^106^ were used to generate a ‘hints’ file for Augustus and GeneMark-ET. Augustus models with at least 90% exon overlap with protein alignments were selected as high-quality models (HiQ). Gene models from the above predictors, HiQ models, transcripts, and predicted proteins were then used as sources of evidence to create consensus models in EVidenceModeler v1.1.1^107^ with weights Augustus:1, HiQ:1, GeneMark-ET:1, PASA:10, SNAP:1, glimmerHMM:1, proteins:1, transcripts:1. The resulting consensus gene models were filtered to remove any proteins less than 50 amino acids, models that began or ended in assembly gaps, and transposable elements. Next, tRNA gene models were identified with tRNAscan-SE v2.0^108^. Final consensus gene models were constructed by funannotate’s ‘update’ function which added and updated UTRs and refined exon boundaries of alternatively spliced models.

To assemble the *de novo* transcriptome for *P. oleracea*, we again filtered all paired-end raw RNAseq reads by quality (phred < 25) and trimmed to remove adapters, poly-A/T tails longer than 12 bp, and low-quality tails (phred < 20) using PRINSEQ v0.20.4. As with *P. amilis*, we pooled all *P. oleracea* RNAseq reads across time points, conditions, and replicates. Pooled reads were input into Trinity v2.11.0 and assembled *de novo*. We reduced the redundancy of the resulting contigs using CD-HIT v4.8.1^109^ by collapsing contigs with at least 98% similarity (-c 98) and open reading frames were identified with TransDecoder. Orthology between *P. amilis* and *P. oleracea* peptide sequences was established using OrthoFinder^110^ v2.5.2 and the primary peptide sequences from genomes of *Ananas comosus*, *Amaranthus hypochondriacus*, *Arabidopsis thaliana*, *Brachypodium distachyon*, *Helianthus annuus*, *Kalanchoë fedtschenkoi*, *Phalaenopsis equistris*, *Sedum album*, *Sorghum bicolor*, *Vitis vinifera*, and *Zea mays*.

### RNAseq differential abundance analysis

We used sleuth^111^ v0.30.0 in R v3.6.1 to assess differential abundance of RNAseq reads quantified by Kallisto for each species independently. For *P. amilis*, we mapped reads to the predicted mRNA transcripts from the genome annotation, and for *P. oleracea* we used the redundancy reduced *de novo* transcriptome. All analyses were conducted at the gene level by providing a transcript-to-gene map to sleuth generated from the final set of gene models and setting ‘gene_mode’ = TRUE. Abundance data were quantified in transcripts per million (TPM) and differential abundance was assessed using a likelihood ratio test between the reduced design matrix, where abundance was purely a function of sampling time (‘y ∼ time_point’), and the full design matrix that included time and water treatment (‘y ∼ time_point + treatment’). *P*-values for unique transcripts were aggregated for each gene model and transformed into *q*-values by adjusting for false discovery rate (Benjamini Hochberg correction).

### Co-expression network analysis

We used Weighted Gene Correlation Network Analysis (WGCNA)^112^ v1.69 in R v3.6.1 to construct co-expression networks and identify modules, and NetworkX^113^ v2.5 in Python v3.7.3 to visualize and measure network features. WGCNA was run separately for each species and condition, resulting in four networks. Using the normalized abundance data for all biological replicates from sleuth, we first divided samples into well-watered and drought conditions for separate network construction (hereafter referred to as the well-watered and drought datasets, networks, modules, etc). We then removed genes from each dataset with maximum expression less than 5 TPM. Remaining genes with excessive missing data or too low variance were removed with the WGCNA function ‘goodSampleGenes’. Outlier samples were detected using hierarchical clustering with ‘hclust’ (stats v3.6.2; R-core team), and removed from each dataset. We determined the soft threshold powers for each network separately using ‘pickSoftThreshold’; the soft threshold power was 8 for both well-watered and drought *P. amilis* networks, and 14 and 7 for *P. oleracea* well-watered and drought networks, respectively. We constructed networks and identified modules using ‘blockwiseModules’ with corType=“bicor”, maxPOutliers=0.05, TOMType=“signed”, deepSplit=2, and mergeCutHeight=0.25 for all networks. With corType=“bicor”, we allowed modules to contain both correlated and anti-correlated expression time profiles. We set minModuleSize=30 for *P. amilis* networks and minModuleSize=100 for *P. oleracea* networks.

After filtering and outlier detection within WGCNA, the final *P. amilis* and *P. oleracea* co-expression datasets contained 26,760 genes for 24 samples and 26,337 genes for 22 samples, respectively. Simultaneous network construction and module detection partitioned genes into 41 well-watered and 21 drought *P. amilis* modules and 29 well-watered and 20 drought *P. oleracea* modules (Supplementary Fig. 1a, b), where modules are mutually exclusive clusters of highly correlated genes. In order to extract photosynthetic gene networks (PGNs), we subsetted the resulting adjacency matrices from WGCNA using over 500 (*P. amilis*) and over 600 (*P. oleracea*) manually curated genes with functions broadly related to photosynthesis and removed edges with adjacency < 0.25. This resulted in networks containing 368-466 genes (Supplementary Table 1). We binned these genes into functional categories reflecting major metabolic pathways (e.g., Calvin-Benson cycle, photorespiration, and starch degradation) and enzymatic reactions or functions (e.g., aminotransferase and PEPC regulation). As many enzymes catalyze reversible reactions or function in multiple pathways, these categorizations are intended to reflect the primary role each protein plays and we recognize that alternative categorizations are possible. For example, RuBisCO is the central protein in both the Calvin-Benson cycle and photorespiration, and MDH functions in both the initial carboxylation pathway that generates malate from CO_2_ and in the decarboxylation pathway that liberates CO_2_ from malate in NAD-type C_4_ biochemistry, albeit to a lesser extent.

### Motif identification and enrichment

Putative *cis*-regulatory motif identification and enrichment was done using the MEME Suite^114^ v5.0.2. First, 5’- and 3’UTRs, introns, and 2kb upstream regions were extracted from all *P. amilis* gene models. The MEME tool^115^ was used to identify *de novo* motifs for each co-expression module and region using an e-value threshold of 0.05 and a minimum motif size of 6bp. We downloaded the JASPAR CORE 2018 motif database^116^, supplemented it with the *de novo* motifs identified in Wai et al.^117^ related to CAM, and finally added *de novo P. amilis* motifs after checking for redundancy with Tomtom^118^. We then used AME (--scoring avg --method fisher --hit-lo-fraction 0.25 --evalue-report-threshold 10.0)^119^ with random sets of control sequences to measure motif enrichment in every module-region combination, as well as the preserved genes of the *PPC-1E1c* and C_4_ modules.

## Acknowledgements

We thank C. M. Mason (University of Central Florida) for identification and collection of *P. amilis*, Karolina Heyduk (University of Hawaii) for insightful comments on early drafts of this work, and Christopher Bolick at Yale’s Marsh Botanic Gardens, who’s support made this work possible. This research was funded by the National Science Foundation (IOS-1754662 to EJE and IOS-1708941 to EWG).

## Data availability

Genomic scaffolds, peptide and coding sequences, assembled transcriptome, and annotations (including repeats) for *Portulaca amilis* are available through the Phytozome web portal (https://phytozome-next.jgi.doe.gov). Raw RNAseq data for *P. amilis* and *P. oleracea* are available through the NCBI’s SRA within BioProject PRJNA732408.

## Code availability

Walkthroughs of genome annotation and RNAseq analyses (including differential abundance and co-expression network analyses) and all custom code are available on GitHub at https://github.com/isgilman/Portulaca-amilis-genome and https://github.com/isgilman/Portulaca-coexpression, respectively.

---

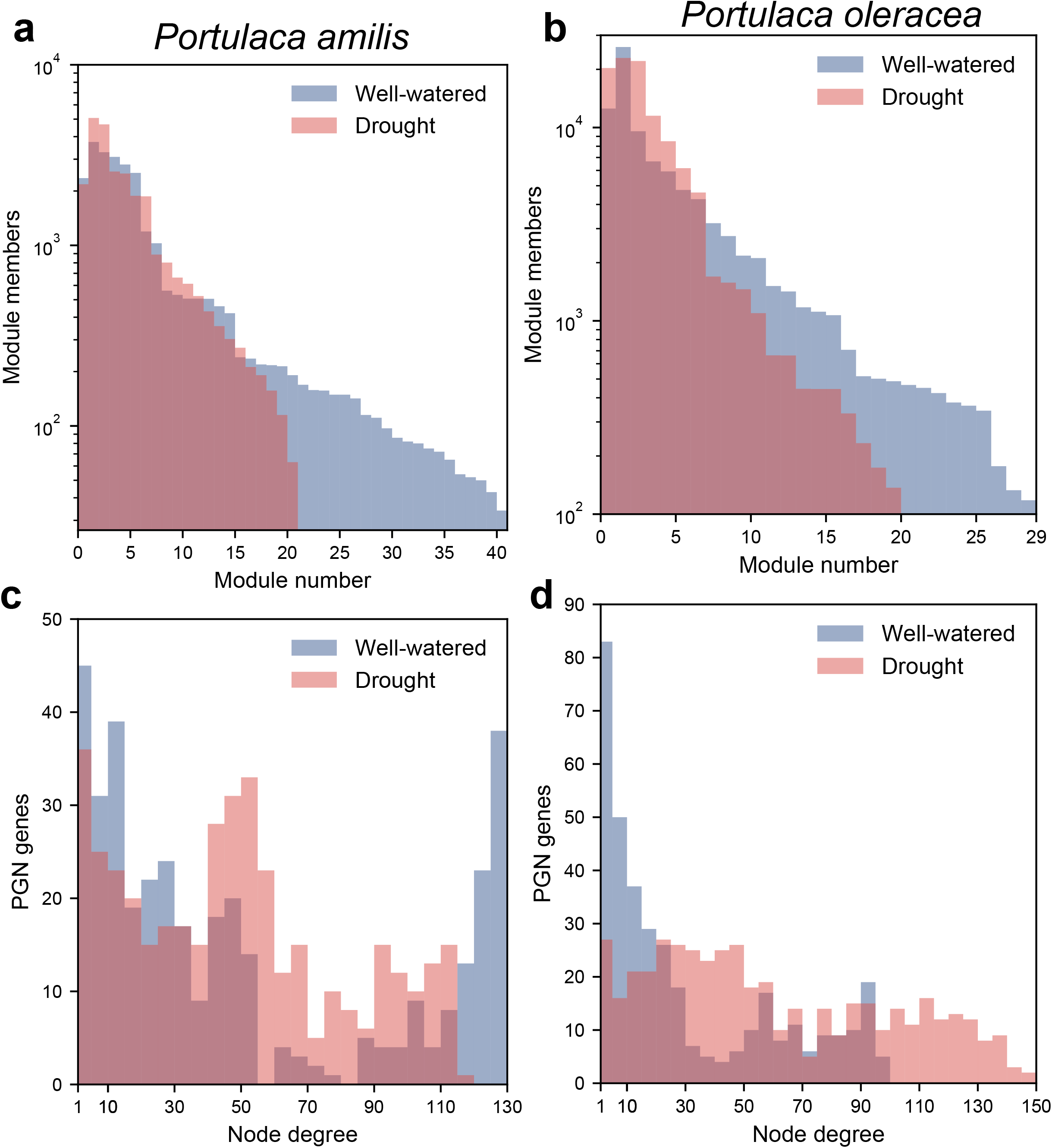

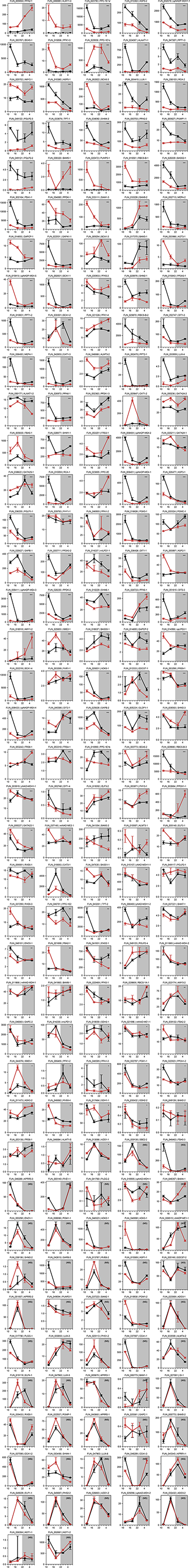

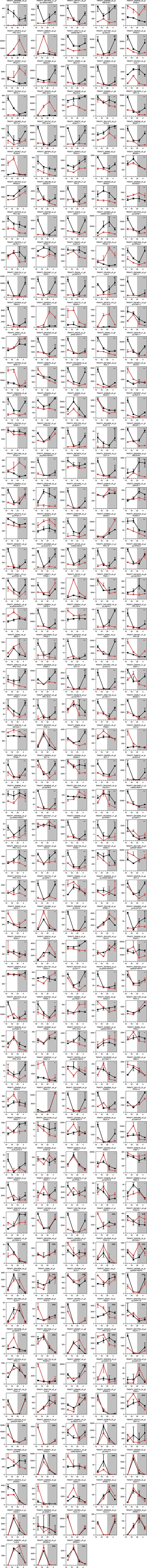

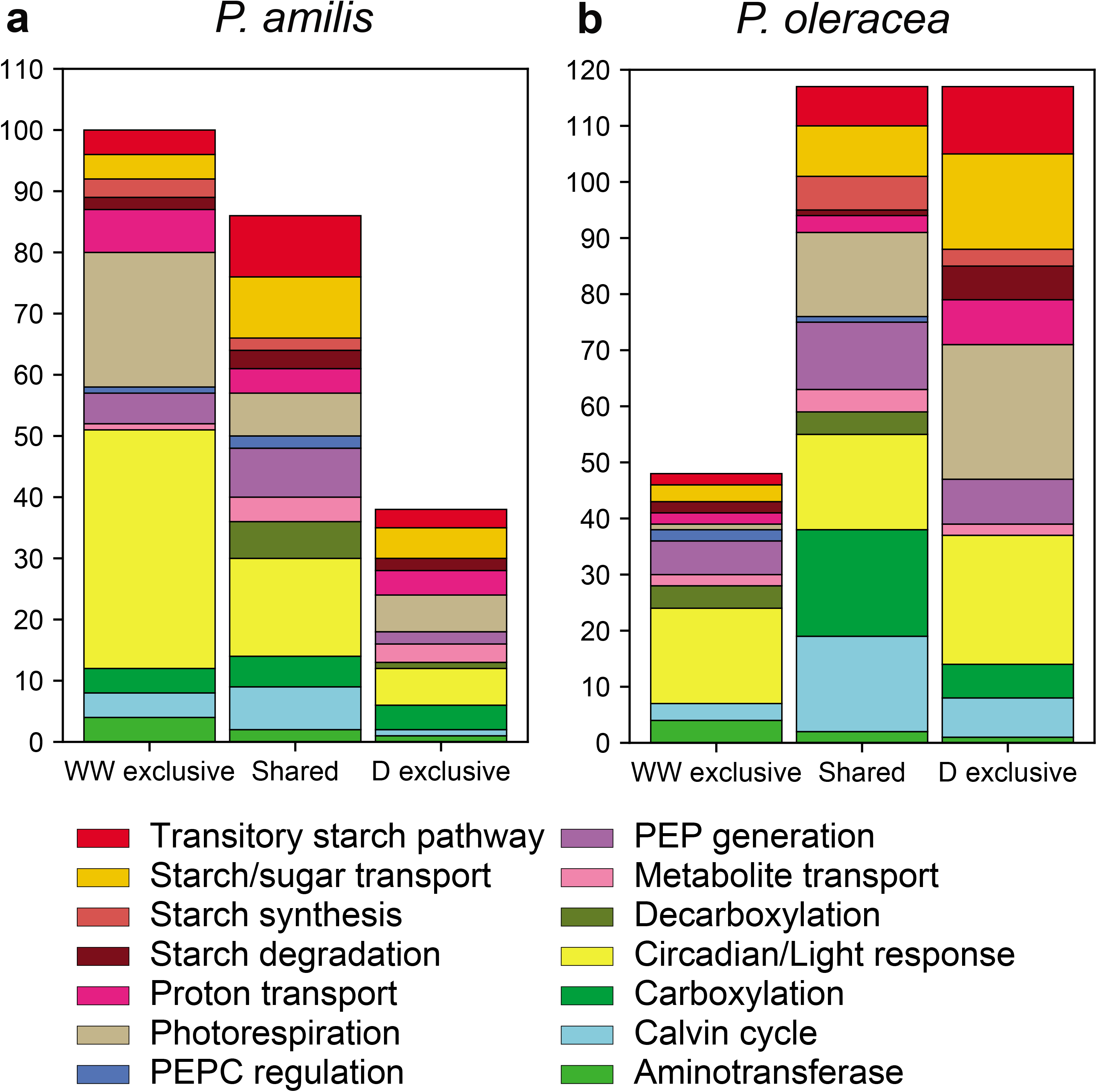

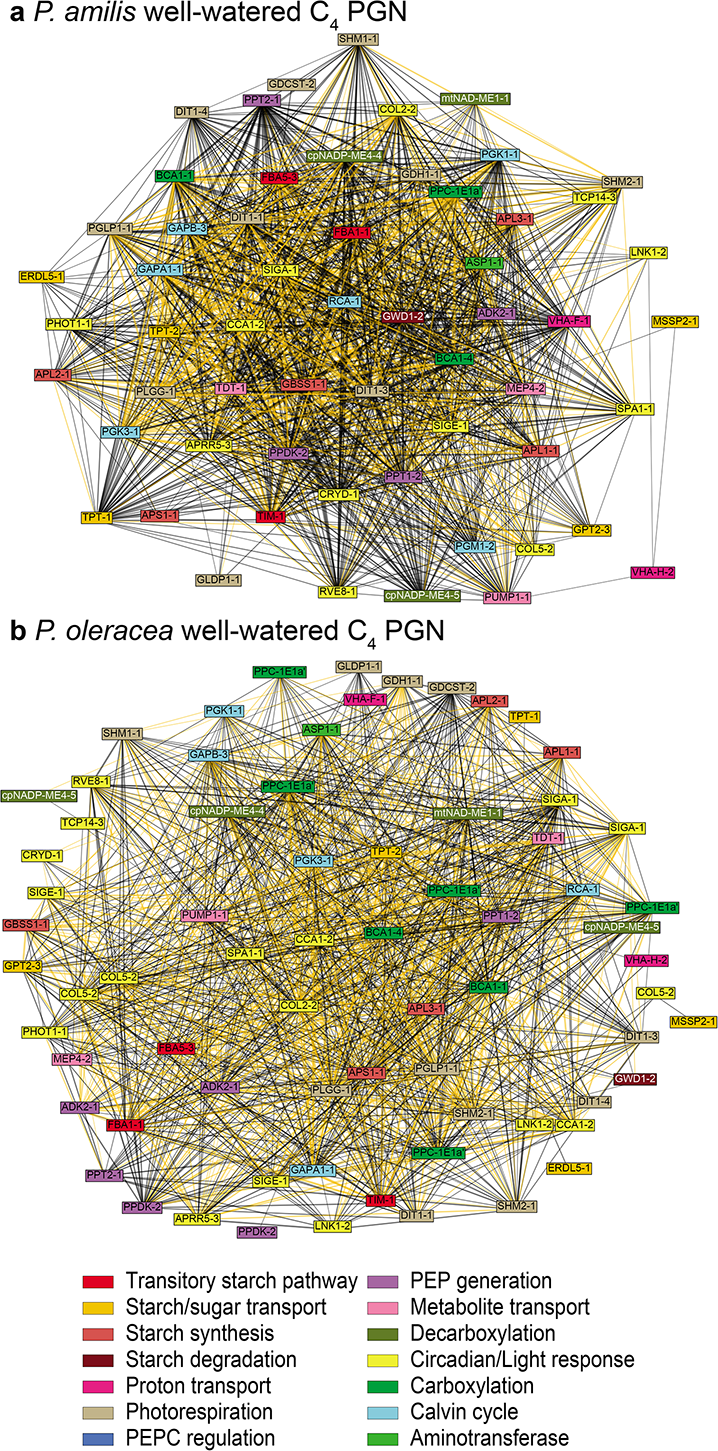

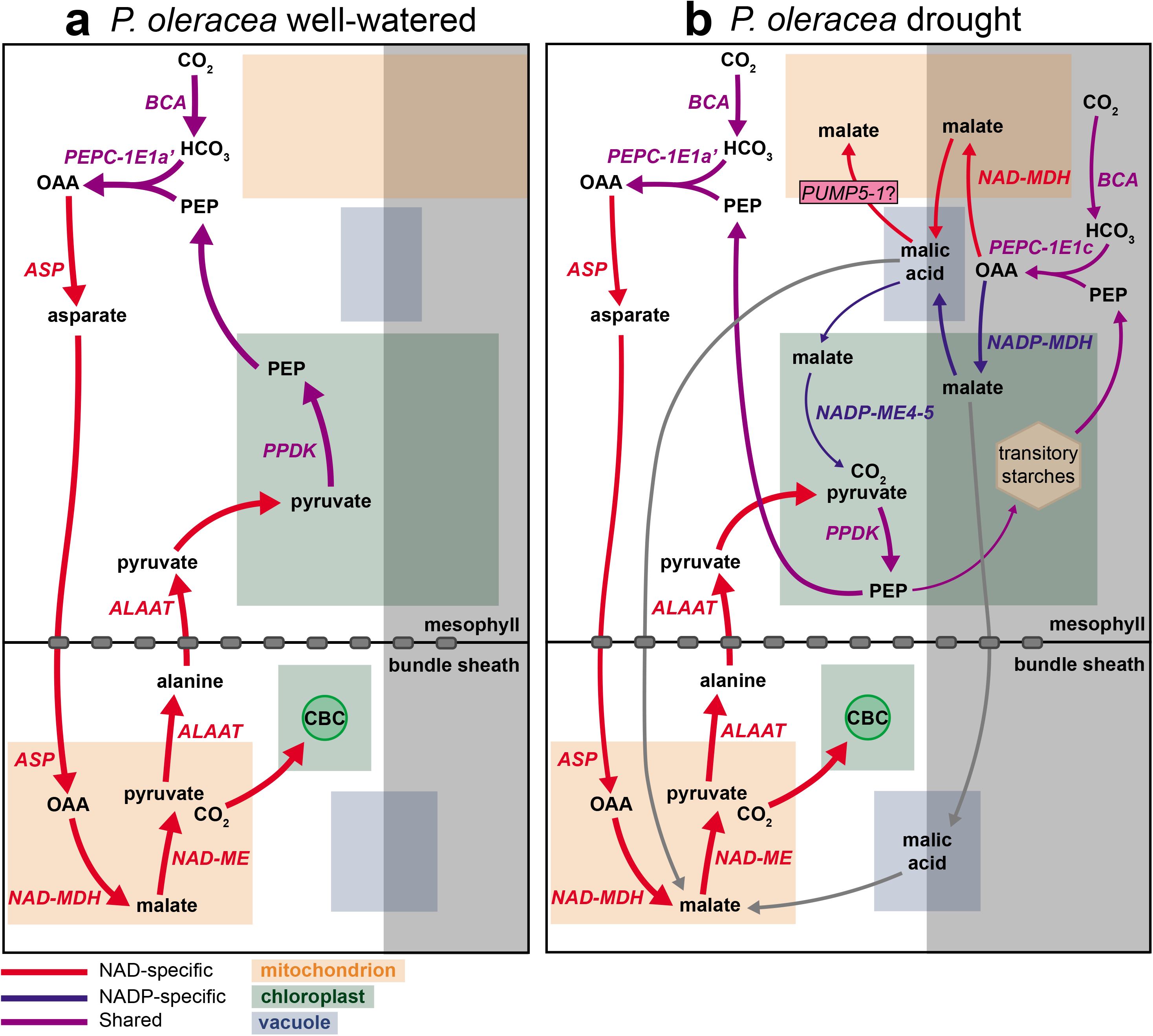

