## Supplementary Text for "Gene co-expression reveals the modularity and integration of C_4_ and CAM in *Portulaca*"

**Supplementary figure legends**

Figure S1. Distribution of module sizes for well-watered (blue) and drought (red) modules of *P. amilis* (A) and *P. oleracea* (B). Histograms of node degree for genes in *P. amilis* (C) and *P. oleracea* (D) photosynthetic gene networks.

Figure S2. Transcripts per million (TPM) normalized abundance of all *P. amilis* gene families discussed in the text. Gene model and name are shown above each plot, and black and red lines indicate median well-watered and drought abundance of all biological replicates, respectively; error bars show interquartile range. Significant differences in expression between well-watered and drought conditions are shown in the upper right corner or each plot; 'NS' = non-significant, '*' = $q < 0.05$, '**' = $q < 0.01$, '***' = $q < 0.001$.

Figure S3. Transcripts per million (TPM) normalized abundance of all *P. oleracea* gene families discussed in the text. Gene model and name are shown above each plot, and black and red lines indicate median well-watered and drought abundance of all biological replicates, respectively; error bars show interquartile range. Significant differences in expression between well-watered and drought conditions are shown in the upper right corner or each plot; 'NS' = non-significant, '*' = $q < 0.05$, '**' = $q < 0.01$, '***' = $q < 0.001$.

Figure S4. Preservation of $C_4$ PGNs in *P. amilis* (A) and *P. oleracea* (B). Orange edges represent correlations among genes shared between species, while black edges are unique to each respective species.

Figure S5. Subnetworks induced by common genes of *P. amilis* (A) and *P. oleracea* (B) well-watered $C_4$ PGNs. Orange edges represent correlations among genes shared between species, while black edges are unique to each respective species. Colors of gene boxes indicate functional categories.

Figure S6. Hypothesized major carbon fluxes of $C_4$+CAM in *P. oleracea* when well-watered (A) and droughted (B). Red, blue, and purple lines indicate NAD-specific, NADP-specific, and shared reactions, respectively, and grey lines show novel, possible pathways for malate that are unique to $C_4$+CAM photosynthesis. Line thicknesses are indicative of relative fluxes through pathways. ALAAT, alanine aminotransferase; ASP, aspartate aminotransferase; BCA, beta carbonic anhydrase; CBC, Calvin-Benson Cycle; NAD-MDH, NAD-dependent malate dehydrogenase; NADP-MDH, NADP-dependent malate dehydrogenase; NAD-ME, NAD-dependent malic enzyme; NADP-ME, NADP-dependent malic enzyme;

PEP, phosphoenolpyruvate; PEPC, PEP carboxylase; PPDK, pyruvate, phosphate dikinase; OAA, oxaloacetate.

**Supplementary tables**

Table S1. Photosynthetic gene network summary statistics

| Network | Nodes (genes) | Edges | Average degree | Median degree | Density * |
|---|---|---|---|---|---|
| *P. amilis* well-watered | 387 | 10,103 | 52.2119 | 34 | 0.1352 |
| *P. amilis* drought | 405 | 9,620 | 47.5062 | 46.0 | 0.1175 |
| *P. oleracea* well-watered | 369 | 5,508 | 29.8537 | 17 | 0.0811 |
| *P. oleracea* drought | 467 | 13,743 | 58.8565 | 49.0 | 0.1263 |

* Density for an undirected graph was calculated by $2m/n(n-1)$, where $m$ is the number of edges and $n$ is the number of nodes.

Table S2. Motifs enriched in the 5'UTRs of genes preserved in the *P. amilis* $C_4$ modules

| Motif name | JASPAR matrix ID | Consensus sequence | Associated TF | *Arabidopsis* ortholog |
|---|---|---|---|---|
| KAN4 | MA1028.1 | GAATATTC | KANADI 4 (KAN4)/AERRANT TESTA SHAPE (ATS) | AT5G42630 |
| OJ1058_F05.8 | MA1033.1 | MCACGTGK | N/A | N/A |
| bZIP68 | MA0968.1 | HNACGTGKM | bZIP transcription factor 68 (BZIP68) | AT1G32150 |

| | | | | |
|---|---|---|---|---|
| ERF122 | MA1261.1 | HDDHTRCGGCKGHGG | Ethylene-responsive transcription factor (ERF112) | AT5G67000 |

**Supplementary data files**

1. *Portulaca amilis* photosynthesis related gene annotations with functional categorizations and module assignments
2. *Portulaca oleracea* photosynthesis related gene annotations with functional categorizations and module assignments
3. Orthogroup assignments
4. Orthogroup gene trees
